# Impact of between-tissue differences on pan-cancer predictions of drug sensitivity

**DOI:** 10.1101/800193

**Authors:** John P. Lloyd, Matthew Soellner, Sofia D. Merajver, Jun Z. Li

**Affiliations:** Department of Human Genetics, University of Michigan, Ann Arbor, MI, 48109, USA; Department of Internal Medicine, University of Michigan, Ann Arbor, MI, 48109, USA; Rogel Cancer Center, University of Michigan, Ann Arbor, MI, 48109, USA

**Author notes:** Co-senior author (Matthew Soellner, Sofia D. Merajver, Jun Z. Li). Co-corresponding author, **Correspondence to:** Jun Z. Li, University of Michigan, Sofia D. Merajver, University of Michigan.

**Keywords:** Cancer biology, MEK inhibitors, drug response predictions, machine learning

## Abstract

Increased availability of drug response and genomics data for many tumor cell lines has accelerated the development of pan-cancer prediction models of drug response. However, it is unclear how much between-tissue differences in drug response and molecular characteristics may contribute to pan-cancer predictions. Also unknown is whether the performance of pan-cancer models could vary by cancer type. Here, we built a series of pan-cancer models using two datasets containing 346 and 504 cell lines with MEK inhibitor (MEKi) response and RNA, SNP, and CNV data, and found that, while the tissue-level drug responses are accurately predicted (between-tissue ρ=0.88-0.98), only 5 of 10 cancer types showed successful within-tissue prediction performance (within-tissue ρ=0.11-0.64). Between-tissue differences make substantial contributions to the performance of pan-cancer MEKi response predictions, as we estimate that exclusion of between-tissue signals leads to a 22% decrease in performance metrics. In practice, joint analysis of multiple cancer types usually has a larger sample size, hence greater power, than for one cancer type; and we observe that the higher accuracy of pan-cancer prediction of MEKi response is almost entirely due to the sample size advantage. Success of pan-cancer prediction reveals how drug response in different cancers may invoke shared regulatory mechanisms despite tissue-specific routes of oncogenesis, yet predictions in different cancer types require flexible incorporation of between-cancer and within-cancer signals. As most datasets in genome sciences contain multiple levels of heterogeneity, careful parsing of group characteristics and within-group, individual variation is essential when making robust inference.

## INTRODUCTION

Tailoring cancer treatment to the molecular characteristics of individual tumors represents one of the central strategies of precision oncology [1–3]. Public repositories of drug response and genomics data in many cancers have facilitated the identification of somatic mutations that underlie variable drug response [3,4]; and drug response prediction analyses have expanded from analyzing known cancer genes to unbiased searches across the human genome [4,5]. More recently, prediction analyses have further incorporated data modalities beyond DNA, to include gene expression, epigenomic, and/or metabolomics data [4,6,7]. Drug response predictions using the tumors’ molecular characteristics are frequently performed within a single cancer type, using primary tumors or auxiliary models derived from them, which are generally expected to share the primary tumors’ vulnerabilities to specific anti-cancer agents – presumably as a result of tissue-specific oncogenic processes involving similar cell types. Meanwhile, drug response predictions can also be developed by considering multiple cancer types jointly, through *pan-cancer* analyses. The success of pan-cancer prediction models is predicated on diverse cancer types sharing a broad set of molecular vulnerabilities despite tissue-specific mechanisms of cancer initiation, progression, or drug-response.

Pharmacogenomic databases of patient-derived cancer cell lines now cover a broad range of cancer types and represent reusable pre-clinical models for finding the cell lines’ innate characteristics that may contribute to their drug response profiles [8–12]. Using tumor cell line resources, many groups have employed DNA variants and gene expression levels to develop pan-cancer drug-response prediction models via a variety of computational methods, including regularized regression [8,13], random forests [14,15], neural networks [16], network modeling [17,18], quantitative structure-activity relationship analysis [19], and deep learning [20]. These analyses have offered many insights, including the importance of RNA expression for pan-cancer predictions [14], higher accuracy of multi-gene classifiers (i.e. gene panels) compared to single-gene classifiers [15], and the suitability of cell lines as *in vitro* mimics of primary tumors [13,14].

Despite such progress, it remains unclear whether between-tissue differences could have contributed to the apparent prediction performance. It is also unclear if the prediction performance could vary among cancer types, a situation that would call for tissue-specific guidelines of applying prediction models. This concern stems from the observation that some cancer types tend to be more sensitive to a certain drug than others, and their tissue-specific molecular properties may drive the performance of a pan-cancer model without necessarily accounting for inter-tumor differences within a cancer type. In clinical practice, while therapeutic decisions are often made solely based on cancer type, there is often the additional need, and potential benefit, to predict variable response among tumors within a cancer type. We set out to examine the relative importance of between- and within-cancer type signals in pan-cancer drug response prediction models. Here, we analyzed data from two public cell line-based datasets [11,12], focusing on ten cancer types that were well-represented in both, and examined the performance of between-tissue and within-tissue models of pan-cancer drug response predictions for MEK inhibitors. In addition to cross-tissue effects, we evaluated cross-MEK inhibitor prediction models, and provided *in silico* replication by applying prediction models across datasets. Based on our results, we highlight key considerations for deploying pan-cancer drug response prediction frameworks and discuss the importance of jointly analyzing the contributions of both group and individual identity when interpreting the performance of prediction models.

## RESULTS

### MEK inhibitor sensitivity across cancer types

To evaluate pan-cancer drug response predictions in pre-clinical tumor models, we utilized publicly available datasets of tumor cell lines described in a prior publication (referred to as Klijn 2015) [11] and the Cancer Cell Line Encyclopedia database (CCLE) [12]. Klijn 2015 and CCLE include 349 and 503 tumor cell lines, respectively, that have drug response data and RNA and DNA characterization, with 154 cell lines in common between the two datasets (**Figure 1A**), representing 10 cancer types defined by organ site (**Figure 1B**). Among the 5 and 24 drugs screened in Klijn 2015 and CCLE datasets, respectively, MEK inhibition was the sole target mechanism in common, with one MEK inhibitor (MEKi) screened in both datasets (PD-0325901; referred to as PD-901) and an additional MEKi unique to each dataset (Klijn 2015: GDC-0973; CCLE: Selumetinib). We therefore focused on predicting MEKi response, as prediction models could be evaluated for consistency across these two independent datasets. Moreover, MEK inhibition has shown promise for pan-cancer drug response predictions [14].

**Figure 1.**
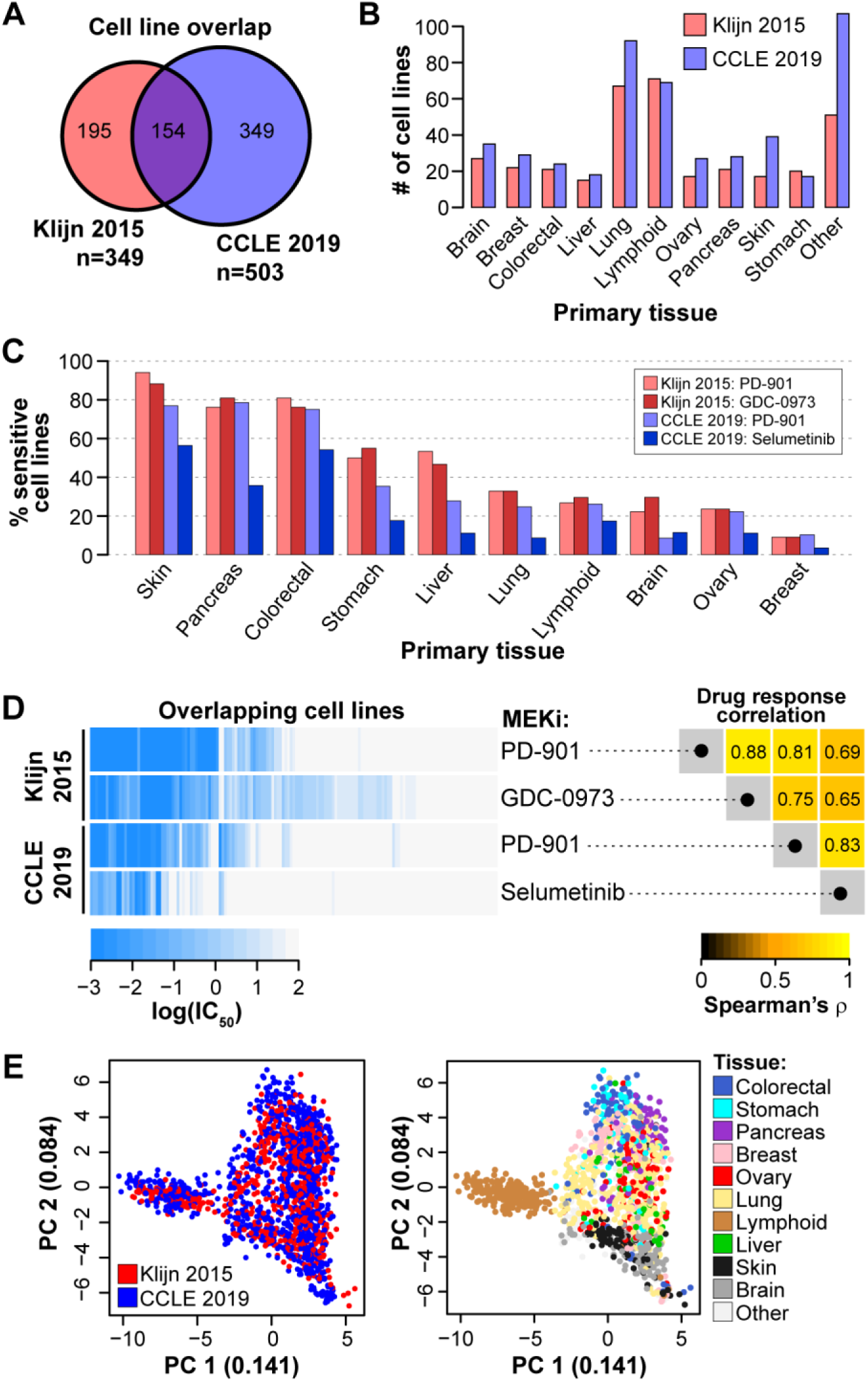
Overview of between-cancer type differences of MEK inhibitor response and expression patterns in the Klijn 2015 and Cancer Cell Line Encyclopedia datasets. **(A)** Overlap in cell lines with drug response and RNA expression and DNA variant data. **(B)** Counts of tumor cell lines with MEKi response data for 10 shared cancer types (n ≥15 in both datasets). **(C)** Proportion of MEKi-sensitive cell lines stratified by tissue. Cell lines were considered sensitive based on a threshold of IC_50_ ≤ 1 nM. **(D)** Variability in the response to the MEK small molecule inhibitors. Left: Heatmap of MEKi response in log(IC_50_) for four data series (in rows). Each column corresponds to a single cell line with drug response data for both MEKi in both datasets; columns were hierarchically clustered (dendrogram not shown). Right: Rank correlation of log(IC_50_) among the four data series in the left panel. **(E)** Scatterplots of the first two dimensions from principal component (PC) analysis on transcriptome data. Each point represents a cell line and is colored by dataset (left) and tissue-of-origin (right).

The molecular features of tumor cell lines available from the Klijn 2015 and CCLE datasets included treatment-naïve RNA expression levels (data from RNA-seq), single-nucleotide polymorphisms (SNPs; Klijn 2015: SNP microarray; CCLE: exome-seq), and copy number variation status for genes (CNVs; SNP microarray). RNA expression data were filtered to include only the 29,279 genes in common between the two datasets. We mapped SNP features to annotated genes, and determined if a gene contained a missense or nonsense SNP (binary variable for the gene: mutated vs. not mutated), while CNV status indicated if a gene was duplicated or deleted (3-level categorical variable: amplified, deleted, or no CNV). SNPs and CNVs had been filtered to exclude known germline variants based on the Exome Aggregation Consortium database [21], but likely still contained rare germline mutations alongside somatic mutations. In all, our DNA data contains mutation status (mutated / not mutated) for 12,399 genes and CNV-carrier status (amplified / deleted / no CNV) for 4,578 genes.

First, we compared MEKi responses of individual cell lines across cancer types and found clear differences. Skin, pancreatic, and colorectal cell lines were generally sensitive to MEK inhibition; while lymphoid, brain, ovary, and breast cell lines were generally resistant (**Figure 1C**). Still, other cancer types, such as the lung, stomach, and liver cancer lines, contain more mixed responses to MEKi (**Figure 1C**). For cancer types with mixed response, i.e. a wide range of observed responses within a cancer type, it would be especially challenging to make treatment decisions based on cancer type alone, hence there is enormous value and opportunity to develop individualized predictions. We further compared the response among the four data series: for two inhibitors in each of two data sources. For the 154 cell lines in common, MEKi response was highly correlated across distinct MEKi and between datasets (Spearman’s ρ=0.65 to 0.88; **Figure 1D**). The observation that cell lines respond similarly to different MEK inhibitors indicates cross-MEKi predictions are feasible, provided the two compounds are chemically similar.

We also compared gene expression profiles between cancer types in each of the two datasets by using principal component analysis (PCA) of standardized RNA expression levels (see Methods; **Figure 1E**). Cell lines from Klijn 2015 and CCLE were jointly analyzed, and they occupy overlapping space in the PC1-PC2 plot (**Figure 1E**). The concordance between datasets supports our approach to develop and test prediction models across datasets (see below). Cell lines from the same primary tissue tend to be present in similar regions of the PC1-PC2 space (**Figure 1E**) and have correlated RNA expression levels (**Figure S1**), highlighting that cell lines derived from the same primary tissue have similar transcriptomic features. Given the effects of primary tissue on both MEKi response (**Figure 1C**) and transcriptomic profiles (**Figure 1E; Figure S1**), it is plausible that between-tissue differences will be a major contributing factor in pan-cancer drug response predictions that consider RNA data, even in cases where tissue labels are not included in a prediction model. The Klijn 2015 and CCLE datasets provide the needed resources for us to investigate the relative importance of between-tissue and within-tissue signals in pan-cancer prediction models.

### Pan-cancer machine learning predictions of MEK inhibitor sensitivity

To build prediction models, we examined two ways of using the drug response variables: either taking log(IC_50_) as a continuous variable or dichotomizing drug response to a binary variable by categorizing cell lines as sensitive (IC_50_ ≤ 1μM) or resistant (IC_50_ > 1 μM). For the former (continuous response variable), we evaluated two prediction algorithms: regularized regression and random forest regression. And for the latter (dichotomized response variable), we adopted two algorithms: logistic regression and binary random forest. As there are two MEKi in two datasets, for each of the four algorithms we developed four prediction models, named as *f_K1_* and *f_K2_* for the two inhibitors in Klijn, and likewise, *f_C1_* and *f_C2_* for the two inhibitors in CCLE. Only the cell lines unique to each dataset were used to train the models (n=195 in Klijn 2015; n=349 in CCLE), while the 154 cell lines in common were saved for validation (**Figure 1A**).

To assess performance, we examined two types of validation: 1) within-dataset validation: using the prediction model developed in a dataset (such as *f_K1_*) to obtain predicted responses for the 154 common cell lines *in the same dataset*, and compare to their observed responses; and 2) between-dataset validation, using the model developed in one dataset to predict the responses for all cell lines - both unique and common lines - *in the other dataset*, and compare to their observed responses. Model performance was evaluated with two metrics: rank correlation (Spearman’s ρ) between observed and predicted log(IC_50_) values, and area under the receiver operating characteristic curve (auROC) on the distributions of prediction scores between cell lines categorized as sensitive (IC_50_ ≤ 1μM) and resistant (IC_50_ > 1 μM). **Figure 2A** shows a workflow where a prediction model, named “*f_K1_*”, is trained on the data for PD-901 (*y_K1_*) in the 195 lines unique to Klijn 2015 dataset (black arrows), and applied to predict (1) within dataset: *ŷ_K(K1)_* for the 154 common lines, using features in Klijn 2015, and (2) across dataset: *ŷ_C(K1)_* for all 503 cell lines in CCLE, including 349 unique and 154 common lines (dashed green arrows). Model performance is assessed by comparing the resulting predicted values from the *f_K1_* model to observed MEKi response (Klijn 2015: *y_K1_* and *y_K2_*; CCLE: *y_C1_* and *y_C2_*; blue dotted arrows). In total, four sets of models { *f_K1_*, *f_K2_*, *f_C1_*, *f_C2_* } were developed using four algorithms each (regularized, logistic, and regression and classification random forest), with predicted MEKi response compared to four observed MEKi response series { *y_K1_*, *y_K2_*, *y_C1_*, *y_C2_*}, with performance measured by two performance metrics (Spearman’s ρ and auROC) for 128 pan-cancer performance measures (**Figure 2B**). As two examples, **Figure 2C** showed the observed and predicted log(IC_50_) values for a within-dataset (regularized algorithm) prediction and a cross-dataset (logistic) prediction from the *f_K1_* model.

**Figure 2.**
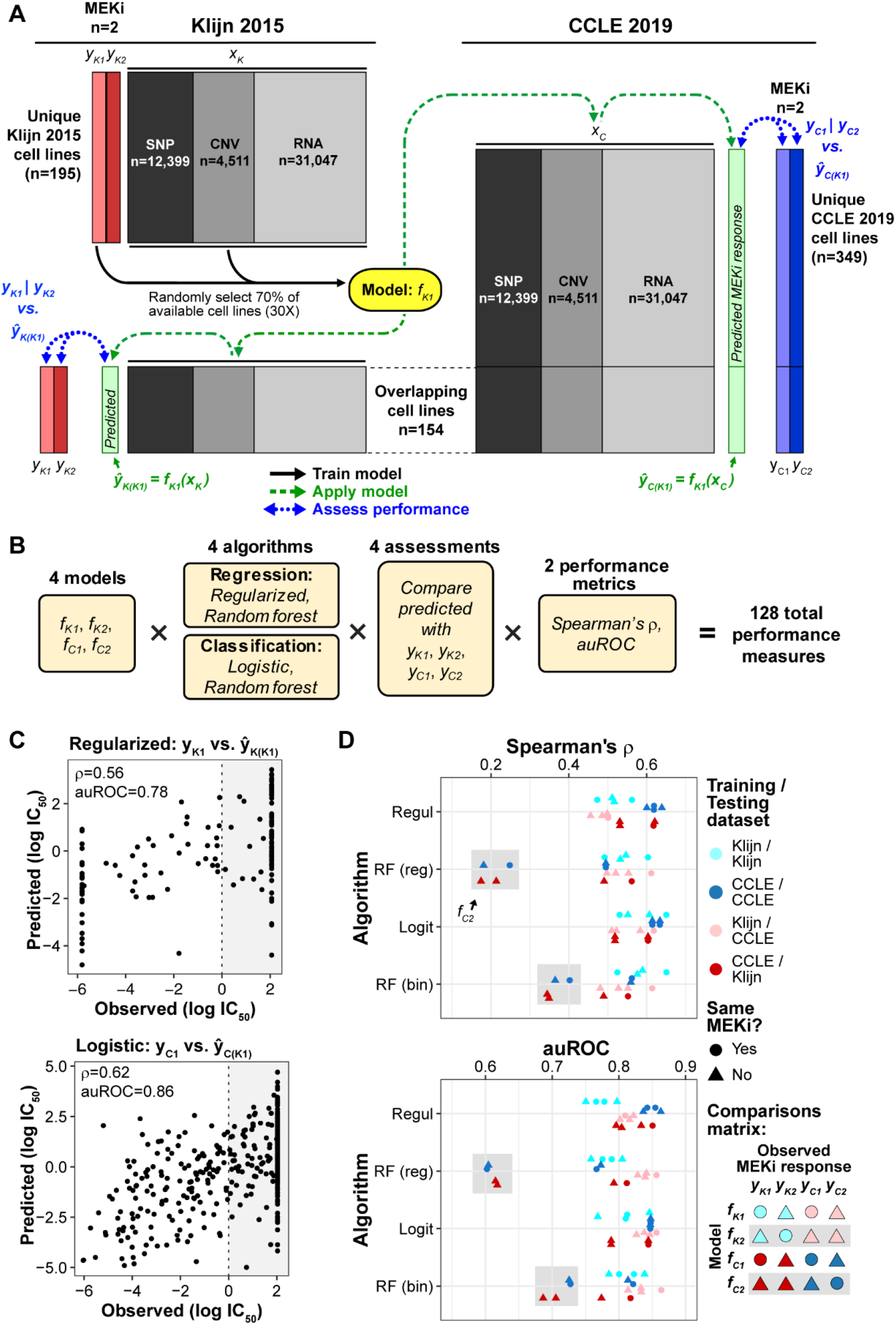
Pan-cancer machine learning predictions of MEKi response. **(A)** Schematic of an example of training-prediction-assessment workflow, depicting the generation of a prediction model (yellow, *f_K1_*) that considers MEKi PD-901 response data from the Klijn 2015 dataset (*y_K1_*, light red) and DNA and RNA features (*x_K_*). The 154 cell lines in common between the two datasets were excluded from model building. Prediction models were built on 70% of training cell lines (selected randomly), repeated 30 times, and the final predicted drug response of a given cell line in the validation sets was calculated as the average of the 30 repeats. The resulting prediction models are applied to within-dataset and cross-dataset RNA and DNA data (*x_K_* and *x_C_*) to generate predicted drug response scores (*ŷ_K(K1)_* and*ŷ_C(K1)_*). Predicted drug response values, shown in light green boxes, were then compared with observed drug response to evaluate model performance (within-dataset: *ŷ_K(K1)_* vs. *y_K2_* | *y_K2_*; cross-dataset: *ŷ_C(K1)_* vs. *y_C1_* | *y_C2_*. Model generation is depicted with black arrows, model application with green dashed arrows, and performance assessment with blue dotted arrows. **(B)** Outline of the full combinations of 4 models based on input data, 4 algorithms, assessments by comparing predicted MEKi response to the 4 series of observed response data, and 2 performance metrics. **(C)** Two examples showing observed and predicted log(IC_50_) from the *f_K1_* model: regularized regression and within-dataset validation (top panel) or logistic regression and cross-dataset validation (bottom). auROC was calculated based on the predicted log(IC_50_) values among cell lines classified as sensitive (log(IC_50_) < 0) and resistant (log(IC_50_) > 0; gray background) regardless of algorithm. **(D)** Performance of all combinations of models, algorithms (y-axis), and assessments by rank correlation (Spearman’s ρ, top panel) and auROC (bottom). Within-dataset performances are indicated by shades of blue: cyan/dark blue, while between-dataset performances are indicated by shades of red: pink/dark red. Models trained from CCLE data are indicated by the darker shade. **Gray boxes**: random forest models trained on CCLE-Selumetinib data (*f_C2_*). **Regul**: regularized regression; **RF (reg):** regression-based random forest; **Logit:** logistic regression; **RF (bin):** classification-based (binary) random forest.

The 128 prediction performances are shown in Figure 2D, with the 4 algorithms arranged in different rows. For regularized and logistic regression algorithms, Spearman’s ρ values ranged from 0.43 to 0.65, and auROC values from 0.74 to 0.86, with similar performance between the two algorithms across different training and testing data (Mann Whitney U test, all *p* ≥ 0.86). Random forest (RF)-based models (both regression and binary) exhibited performance similar to the regularized and logistic regression models, except for the *f_C2_* models: those trained on Selumetinib response and RNA and DNA features in CCLE, which exhibited lower performance relative to other algorithm × training set combinations (gray boxes in **Figure 2D**). In the following sections, we focus on predictions from the regularized regression and logistic regression algorithms, as these algorithms are simpler and performed similarly or better than the more complex random forest algorithms.

We next compared the performance from the perspective of cross-dataset and cross-MEKi predictions. Prediction models applied across datasets (red and pink symbols in **Figure 2D**) performed similarly to those applied within datasets (blue and cyan symbols) for regularized and logistic regression algorithms, based on either Spearman’s ρ or auROC (U test, all *p* ≥ 0.06; **Figure 2D**). Similarly, we found that within-MEKi (circles) and cross-MEKi (triangles) predictions performed comparably (U test, all *p* ≥ 0.11; **Figure 2D**). Overall, our results are consistent with those previously reported that response to anti-cancer drugs can be successfully predicted based on pan-cancer datasets [8,14,15,20,22]. Importantly, our framework provided strong *in silico* validation of the models’ accurate cross-dataset predictions, as MEKi response and RNA-DNA data in the Klijn 2015 and CCLE datasets were collected independently.

### Between- and within-tissue performance of pan-cancer MEKi response predictions

To evaluate between-tissue performance, we averaged the observed and predicted values for cell lines within each cancer type and asked if the model recapitulated the group-level differences across the 10 cancer types. The mean observed and mean predicted log(IC_50_) values were highly correlated across cancer types (ρ range: 0.88–0.98, all *p* < 0.002; four comparisons shown in **Figure 3**, mean values as ellipsis centers), indicating that the pan-cancer prediction models accurately captured the average drug responses of tissues. Next, to evaluate within-tissue performance, we calculated the correlation between the observed and predicted log(IC_50_) values for cell lines *within* a single cancer type, and depicted the strength of correlation as the shape and tilt of the ellipsoid for individual cancer types (**Figure 3**). Based on hierarchical clustering of the resulting within-tissue prediction performances from different algorithms and training/testing sets (**Figure S2**), we identify five tissues whose within-tissue variability was accurately predicted by pan-cancer prediction models: liver, ovary, lymphoid, stomach, and skin (mean within-tissue ρ = 0.59; ρ range: 0.51-0.64; **Figure S2**). The remaining five tissues (pancreas, brain, colorectal, breast, and lung) exhibited lower within-tissue prediction performance (mean ρ = 0.28; ρ range: 0.11-0.40; **Figure S2**). This result shows that whether a pan-cancer MEKi response prediction model is informative for an individual tumor (or patient) depends on which tissue is considered, with half of the tissues evaluated here achieving within-tissue predictions with ρ ≥ 0.51.

**Figure 3.**
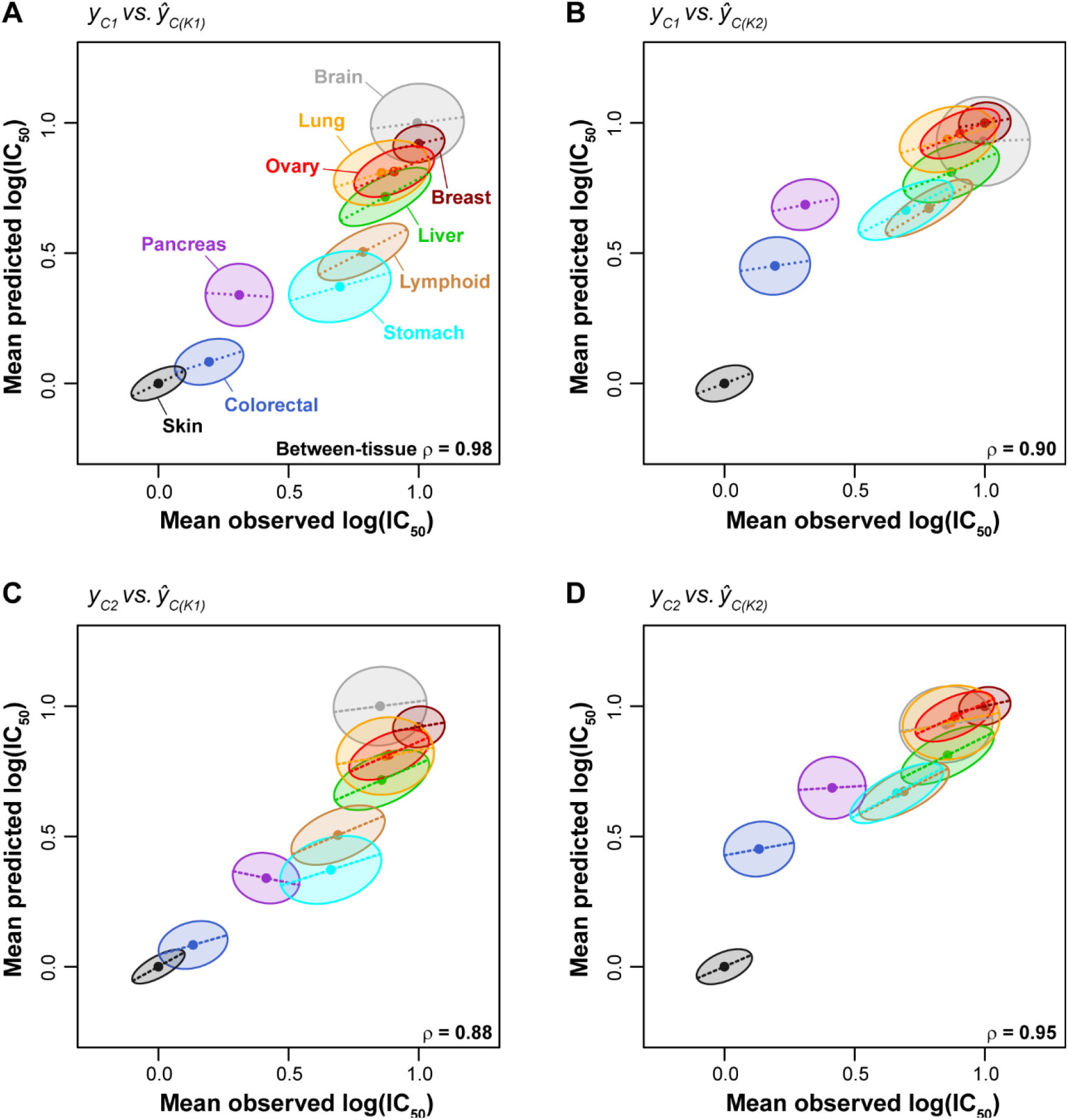
Cigar plots of pan-cancer MEKi response predictions within and between cancer types. Dots at the center of ellipses indicate the mean observed and predicted log(IC_50_) values for a given tissue (mean values were scaled linearly between 0 and 1). ρ values indicate rank correlation among ellipsis centers. Maximum ellipsis length scales with the range of MEKi responses for a given tissue, where tissues with larger response ranges are associated with longer ellipses (e.g. stomach and lymphoid) while tissues with a smaller response range are associated with shorter ellipses (e.g. skin and breast). The slope of dashed lines (and tilt of ellipses) corresponds to the within-tissue regression coefficient of the predicted values against the observed values. The width of ellipses also corresponds to the within-tissue regression coefficient, i.e., a high correlation value is shown as a slender ellipse while a low correlation value leads to a round ellipse. **(A-D)** Within- and between-tissue performances for the 4 combinations of drug/models trained on Klijn 2015 data and applied to CCLE data.

Accurate between-tissue predictions increase the overall prediction performance, which can be seen most clearly if we focus on two of the tissues, the brain and pancreas cell lines. Although within-tissue performance for either brain or pancreas cell lines was poor (both ρ ≤ 0.16 for *y_C1_* vs. *ŷ_C(K1)_*’, **Figure 4A; Figure S2**), prediction performance increased dramatically when the two tissues are considered together (ρ = 0.54; **Figure 4A**). This is because the two tissues, at the group level, are different in both drug response and molecular profiles (**Figure 1C, E**) and the overall performance is largely driven by between-tissue signals. To quantify the contribution of between-tissue signals on the performance of pan-cancer predictions, we standardized the observed and predicted MEKi response within tissues so that the per-tissue mean values are centered (an example in **Figure 4B**), and assessed the change in performance calculated from tissue-standardized values relative to the initial, non-tissue-standardized values (**Figure 4C, D**). Compared to the initial performance, the rank correlation coefficients using tissue-standardized observed and predicted log(IC_50_) values were reduced by an average of 22% (range: 3-35%), depending on the algorithm and training/testing data (**Figure 4C, D**).

**Figure 4.**
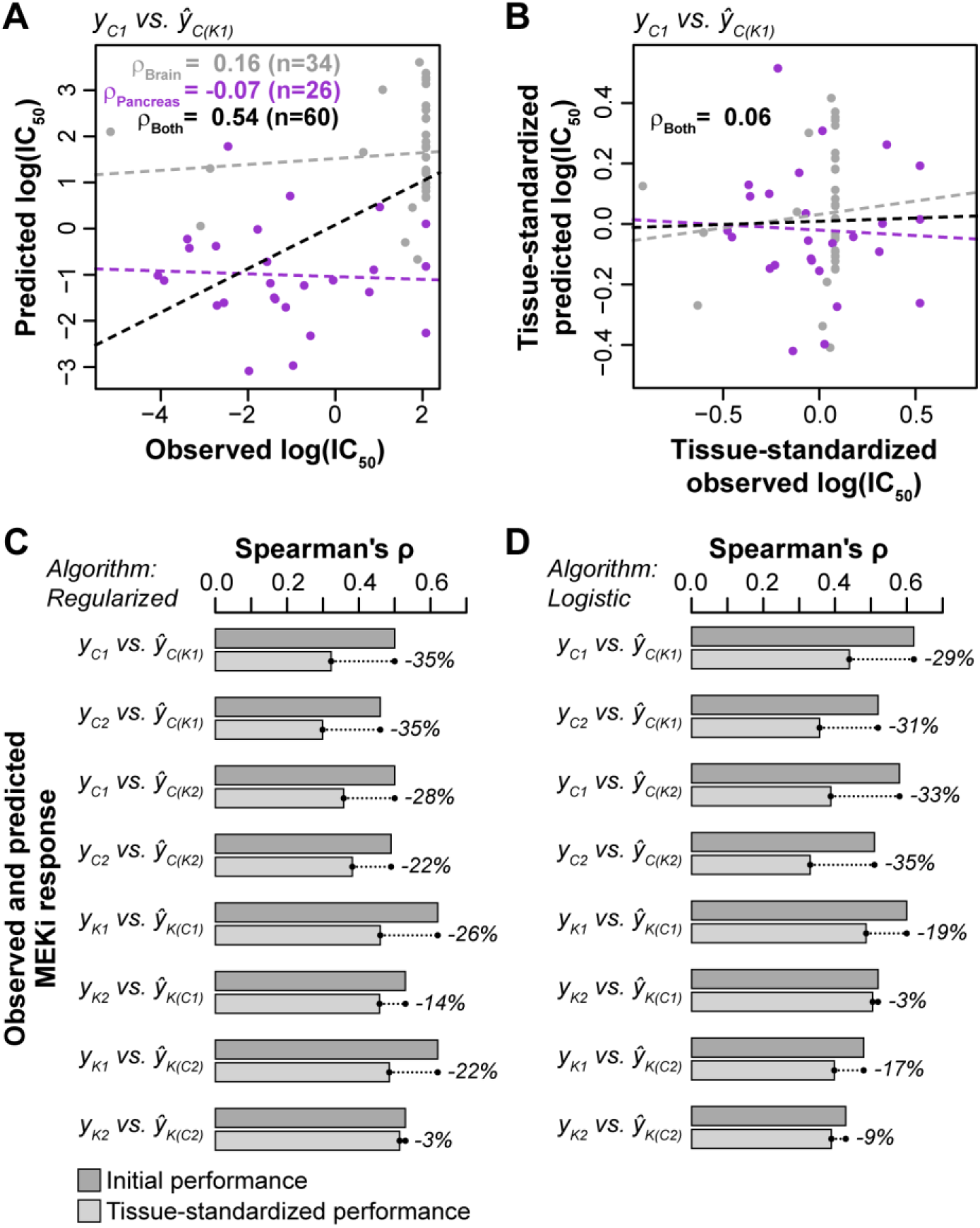
Effects of between-tissue signals on prediction performance. **(A)** Performance of pan-cancer prediction models for a combination of brain and pancreas tissues. Rank correlation (Spearman’s ρ) is shown for brain cell lines (gray), pancreas cell lines (purple), and both brain and pancreas combined (black). Dashed lines indicate lines of best fit. **(B)** Performance of pan-cancer prediction models for a combination of brain and pancreas tissues following standardization of observed and predicted MEKi response within tissues. Standardization was performed by scaling linearly between 0 and 1, followed by subtraction of the scaled mean. **(C, D)** Comparisons of initial performance (dark gray bars; **Figure 2D**) vs. performances calculated following standardization of observed and predicted log(IC_50_) values within all tissues (light gray bars) for **(C)** regularized and **(D)** logistic regression algorithms. Percentage values indicate the % decrease of ρ in the tissue-standardized assessment compared to the initial assessment.

At the feature level, we evaluated the top 50 biomarkers - those having the highest regularized regression weights - for each model, and found that, as expected, they explained a significantly higher amount of within-tissue MEKi response variation (all *p* < 4×10^−11^, U tests; **Figure S3**) than other features. Importantly, they also carry strong between-tissue signals: having significantly higher correlation between mean within-tissue gene expression and mean within-tissue MEKi response across 10 tissues (all *p* < 9×10^−12^, U tests; **Figure S3**). Overall, these results indicate that between-tissue signals make a strong contribution to the performance of pan-cancer drug response predictions and markers most important to the models often contain both within-tissue and between-tissue signals.

### Sample size advantage of pan-cancer models over tissue-specific models

We next asked if pan-cancer prediction models can outperform those generated by considering a single cancer type. To address this question, we compared single-tissue prediction models (i.e. trained and tested with cell lines from a single cancer type) and pan-cancer prediction models. As CCLE has a larger sample size, we considered only regularized regression models trained on CCLE and tested on Klijn 2015. We found that for the liver, lymphoid, and colorectal cancers, the performance of pan-cancer model is similar to that of single-tissue model; whereas for the other six cancer types: ovary, stomach, skin, lung, breast, and brain, the pan-cancer predictions are more accurate (**Figure 5**). In all, the pan-cancer models performed better than or as well as tissue-specific models for 9 out of 10 cancers, whereas pancreas was the only cancer type for which tissue-specific models outperformed the full pan-cancer models.

**Figure 5.**
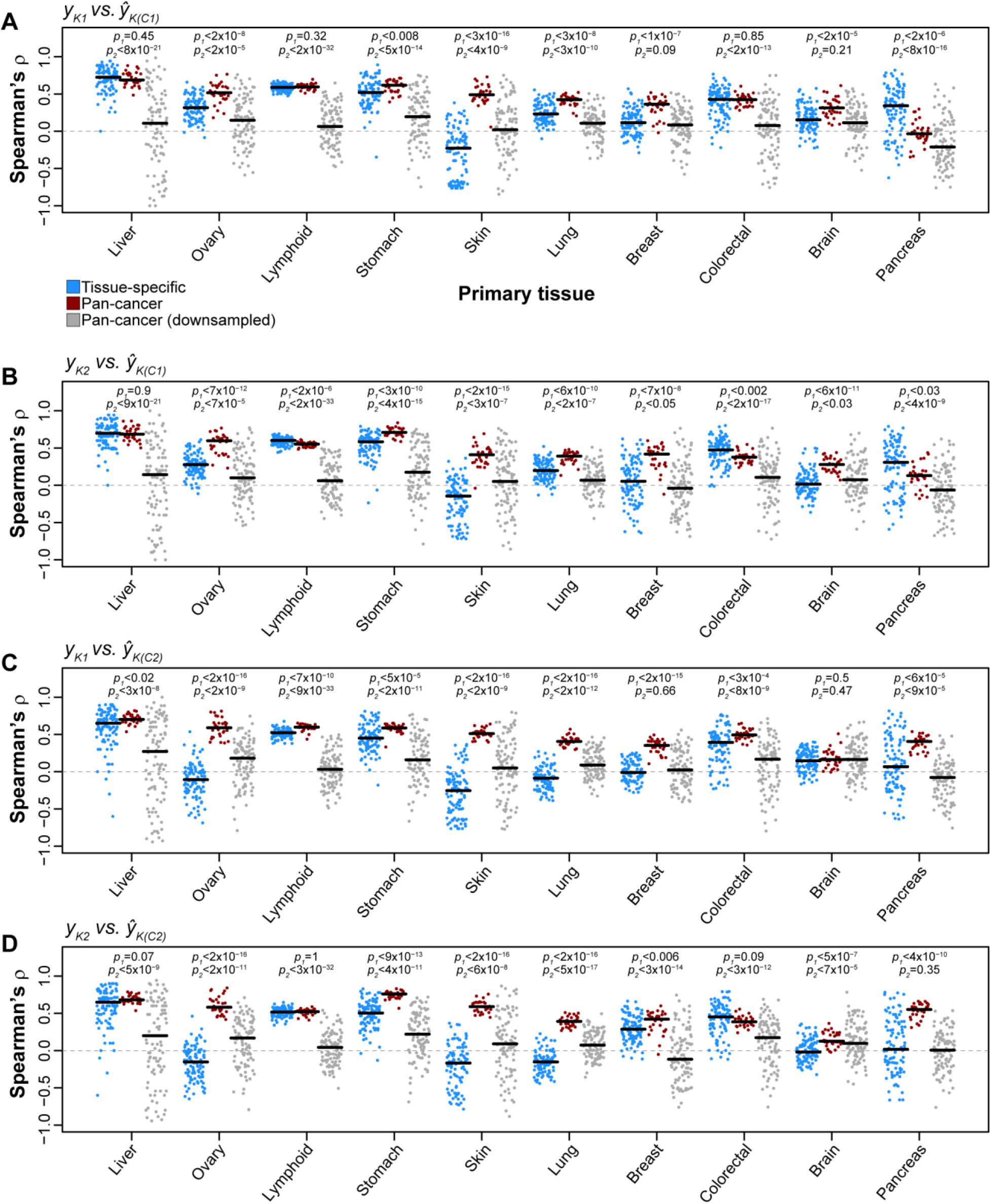
Comparisons of pan-cancer and tissue-specific MEKi response prediction models. For each tissue with ≥15 cell lines in both Klijn 2015 and CCLE datasets, a tissue-specific prediction model was trained and tested by considering only cell lines from a given tissue. Regularized regression prediction models were trained on a random selection of 75% of cell lines in CCLE data of a given tissue type and applied to Klijn 2015 cell lines of the same tissue type (repeated 100 times). Performance was reported using rank correlation between observed and predicted MEKi drug response for each iteration (blue points). Rank correlation for the 30 iterations of MEKi response predictions based on pan-cancer prediction models (**Figure 2**) for a given tissue are indicated with red points. Pan-cancer prediction models were trained and tested using many more cell lines than tissue-specific prediction models. A new set of pan-cancer prediction models were generated by downsampling the available pan-cancer cell line sets to sample sizes equal to tissue-specific prediction models (gray points). **(A-D)** Results are shown for the 4 combinations of drug-models trained using CCLE data and applied to Klijn 2015 data. **Black horizontal lines**: median performance for a given distribution. **p_1_**: P-value from a Mann-Whitney U test of the difference between tissue-specific and pan-cancer prediction performances for a given tissue. **p_2_**: P-value of the difference between tissue-specific and downsampled pan-cancer prediction performances.

A caveat to this result is that pan-cancer models were trained with 4- to 28-times more cell lines than tissue-specific models (e.g. 68 lung and 12 liver cell lines for tissue-specific models). To evaluate the effect of sample size we added an analysis where we down-sampled the pan-cancer data to the same sample size of the single-tissue models (“Pan-cancer (downsampled)” in **Figure 5**). These equal-sized pan-cancer models rarely outperformed tissue-specific models, except in cases where tissue-specific models perform particularly poorly (e.g. skin and lung cancers). Thus, the advantage of pan-cancer prediction models is only apparent when they are based on a larger sample size than tissue-specific models. At comparable sample size, tissue-specific models tend to be more accurate.

### Estimating sample sizes required for optimal prediction performance

The analysis above raises the practical question: how many cell lines are needed to produce optimal performance for pan-cancer models? To assess the relationship between prediction performance and sample size, we developed regularized regression models from a series of randomly downsampled sets of pan-cancer cell lines (n_cell lines_ = 20-300; step size = 10; n_iterations/step_ = 30). For this analysis, we developed only downsampled *pan-cancer* prediction models, and assessed performance in both pan-cancer and tissue-specific testing sets. For the tissue-specific testing subsets, we focused on cell lines from liver, ovary, lymphoid, stomach, and skin tissue, as these five tissues were well-predicted by pan-cancer prediction models (as shown in **Figure 3**; **Figure S2**). For overall pan-cancer prediction, we observed an inflection point for training sample size between 70 and 100 cell lines (~70 cell lines for *f_C1_*, **Figure 6A, B**; ~100 cell lines for *f_C1_*, **Figure 6C, D**). Including additional cell lines further increased performance, but with diminished rate of improvement. Tissue-specific predictions by pan-cancer models with >100 cell lines resulted in diminishing returns for lymphoid and stomach cell lines, while the performance reached saturation even more quickly for liver and skin cell lines (**Figure 6**). However, performance for ovary cell lines showed no inflection point; it continued to increase as additional cell lines were included (**Figure 6**), suggesting ovarian cancers may be well-positioned to benefit from even larger pan-cancer datasets. Overall, we find that generally 70-100 samples are needed to provide robust pan-cancer prediction performance in most cases.

**Figure 6.**
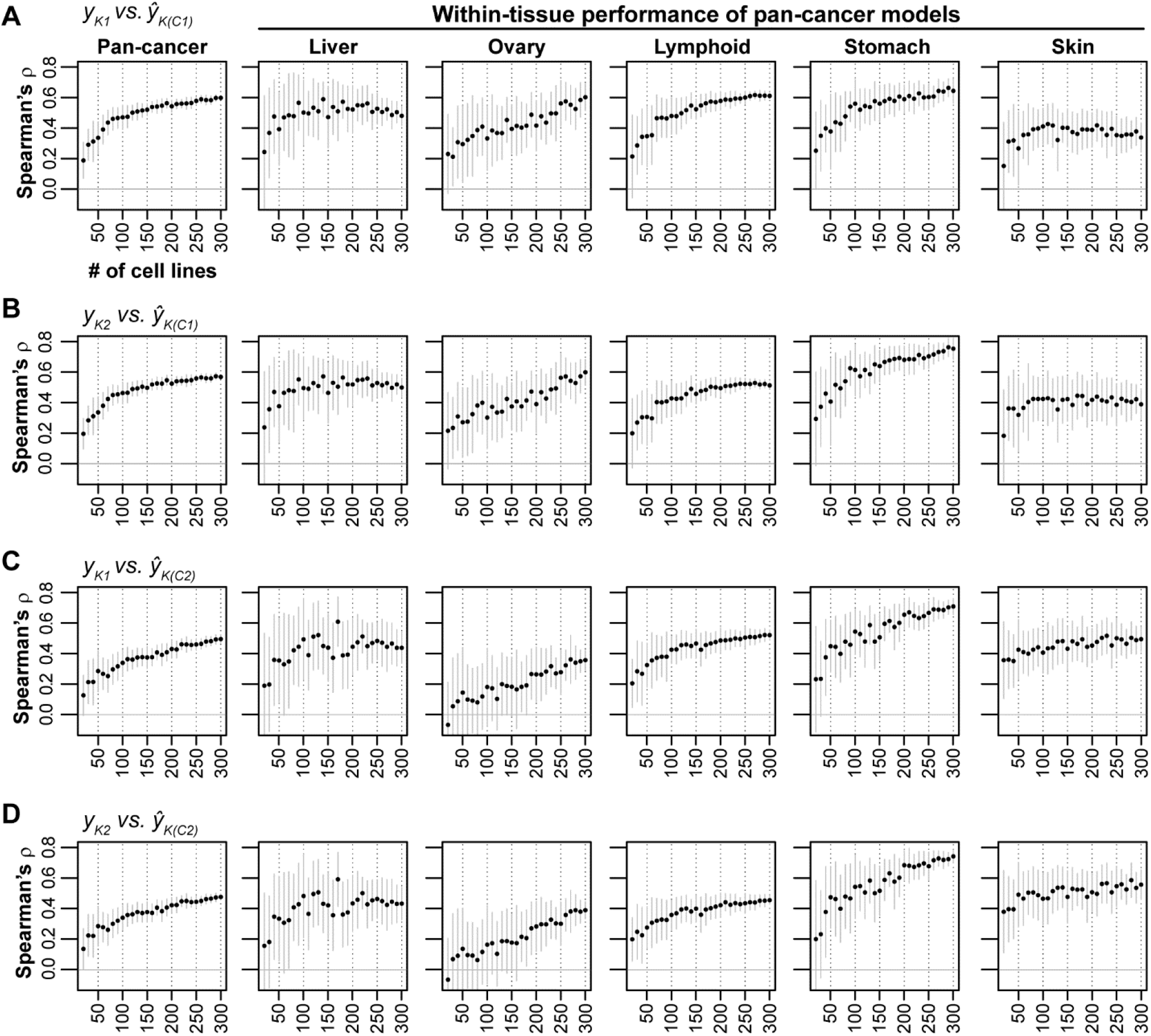
Performance of pan-cancer MEKi response predictions based on downsampled sets of cell lines. Downsampled pancancer models were applied, from left to right, on pan-cancer prediction and five single-tissue predictions. Cell lines in the CCLE dataset were randomly downsampled to sample sizes ranging from 20 to 300 in 10 cell line increments (x-axis; n_iterations_=30). Regularized regression prediction models were trained on downsampled cell lines sets and applied to cell lines in the Klijn 2015 dataset. Mean ranked correlation (Spearman’s ρ) between observed and predicted logged IC_50_ values across the 30 iterations for each sample size are plotted. **(A-D)** Results are shown for the 4 combinations of drug-models trained using CCLE data and applied to Klijn 2015 data. **Solid gray vertical lines**: ±standard deviation.

## DISCUSSION

The feasibility of pan-cancer drug response prediction is predicated on the existence of shared molecular vulnerabilities across cancer types. When successful, these predictions can contribute to the practice of precision oncology by informing individualized treatment decisions. In this study, by using pharmacogenomic datasets for MEK inhibition we confirmed that pan-cancer models did produce the level of performance (**Figure 2C**) reported in similar studies. However, unlike other studies, we underscore the observation that between-tissue differences make a substantial contribution to the apparent performance of pan-cancer models. This is because many cancer types differ in both molecular features and drug response. The effects of between-tissue signals on the prediction models are reflected in the markedly decreased performance when only within-tissue effects are examined (**Figure 4**). Similarly, biomarkers most important to the prediction models tend to carry both within- and between-tissue signals (**Figure S3**). As datasets in genome science typically contain multiple levels of heterogeneity, assessing the contributions of between- and within-group signals in prediction models represents an issue that extends beyond pan-cancer drug response predictions. We also find that pan-cancer models can outperform tissue-specific models, although this advantage usually disappears at comparable sample sizes (**Figure 5**). In practice, when the sample size for an individual cancer type is small, pan-cancer models can be developed from a larger, mixed-type training set, and applied as a powerful tool for informing anti-cancer therapies. The benefit is especially pronounced for the cancer types that contain a wide within-type variation of responses (e.g. MEK inhibition drugs in lung, liver, and stomach cancers).

While our analyses are limited to two publicly available datasets that contain gene expression, SNP, and CNV data, we can draw several lessons. First, the overall prediction performance across all cancer types include both between-tissue and within-tissue effects. Additional assessment and reporting for individual cancer types is recommended. Second, prediction performance is variable between cancers of different tissue origin. As a result, a pan-cancer prediction score should be used with greater caution for an under-performing drug-tissue setting (e.g., MEK inhibitor response in pancreas, brain, or colorectal cancers). In practice, the prediction score for a given tumor may come with a claim of high accuracy if such scores are evaluated in a pan-cancer setting, but for the individual variability within the tumor type it may offer little predictive power. In this situation, using the prediction score would be akin to treating the tumor as an average sample of that tumor type, thus not much different from making a decision based on the tumor type. Third, biomarkers identified using pan-cancer models can emerge for different reasons: some driving a strong response in one of a few tissue, others involved in drug-response programs shared in common by many more tissues, still others could simply be tissue-specific genes not involved in drug-response. Marker selection in the future will need to consider the diversity of tissue origin in the training panel as well as the tissue type of the application.

We also highlight three observations that are relevant to designing future experiments or investigating additional cancer models and drug families. First, when assessing sample size of pan-cancer models for MEKi, we show that inclusion of more than 100 cell lines frequently led to diminishing performance returns. Second, pan-cancer models for one MEK inhibitor (e.g. PD-901) led to successful predictions for other MEK inhibitors (e.g. Selumetinib; **Figure 2**). The MEK inhibitors considered in this study share a specific mechanism of action: potent inhibition of the MEK1 protein (although PD-901 also inhibits MEK2) [23]. Third, across the ten cancer types, although lymphoid cell lines are biologically and clinically distinct from the other, epithelial-origin cell lines (see **Figure 1E**), the pan-cancer models can accurately predict this cancer type (**Figure 3**; **Figure S2**). This suggests lymphoid cell lines may respond to MEK inhibition via similar biological pathways.

With the ever-increasing availability of large pharmacogenomic datasets and the introduction of basket clinical trials, which select patients based on tumor features rather than cancer type, pan-cancer prediction models and biomarker sets will be increasingly adopted to match patients with effective treatments. Dissecting between-tissue and within-tissue statistical signals in drug response prediction models is essential for contextualizing “precision” in precision oncology, allowing the balanced consideration of group-based (i.e. tissue class) treatment decisions on one hand, and individual-based decisions on the other.

Our examination of between- and within-tissue signals in pan-cancer drug response prediction models echoes similar concerns in other fields that involve joint contributions of individual and group factors. For example, shared ancestry is a known confounder in genomewide association studies, where correlations between traits and ancestry can lead to false positives in association tests and mis-estimates of marker effects, as has been demonstrated for variation in anthropomorphic traits (e.g. height) or disease risks (e.g. Type 2 diabetes) [24–28]. Similarly, forensic markers chosen based on their ability to identify individuals also carry population-level information [29]. Plant and animal breeding approaches that utilize genomewide SNP patterns to predict economically-important traits, known as genomic selection, must be adjusted to ensure informative markers explain variation in the trait of interest, rather than population structure [30,31]. Ultimately, developing an understanding of both the group-level and individual-level factors affecting individual trait values is needed in interpreting prediction models and marker panels, especially those generated from highly heterogeneous populations.

## METHODS

### Cell line annotations and drug response data

Drug response and molecular characterization data were retrieved from two sources: 1) the supplemental material of a prior publication (Klijn 2015) [11] and 2) a tumor cell line database (CCLE) [12]. For Klijn 2015, cell line annotations, including identifiers and primary tumor types, were retrieved from “Supplementary Table 1” in [11]. Annotations for CCLE were retrieved from “Supplementary Table 1” in [12]. Cell line identifiers were cross-mapped across datasets and data modalities (i.e. drug response, RNA expression, and DNA variants) by accounting for differential use of upper/lower case characters and/or non-alphanumeric characters; alphabet characters in cell line identifiers were converted to upper case and identifiers were stripped of non-alphanumeric characters. Klijn et al [11] excluded 65 cell lines from analysis due to SNP similarity to other cell lines or uncertainty in cell line origin; we also excluded these cell lines from our analyses (“In Final Analysis Set” column = FALSE; “Supplementary Table 1” in [11]).

For Klijn 2015, we retrieved drug response data from “Supplementary Table 13” in [11]. Drug response was provided for 5 drugs as IC_50_: the dose at which 50% of cells are non-viable. Lower IC_50_ values indicate greater sensitivity. Among the 5 drugs, we focused on two MEK inhibitors: PD-0325901 (referred to as PD-901) and GDC-0973. For CCLE, drug response data were downloaded in the file “CCLE_NP24.2009_Drug_data_2015.02.24.csv” (database file date: 24 February 2015), which included IC_50_ data for 24 anti-cancer drugs. We used drug response data for two MEK inhibitors in CCLE: PD-901 and AZD-6244 (trade name: Selumetinib). The natural log of IC_50_ values were used for feature selection, parameter sweeps, and model training and evaluation.

### RNA expression data

For Klijn 2015, we analyzed expression data from RNA-seq available from two supplemental datasets, designated as protein-coding (“Supplementary Data 1” in [11]) and non-coding (“Supplementary Data 2” in [11]). In all, there were 25,984 coding genes and 21,398 non-coding genes after excluding genes that were invariable in expression across all 675 cell lines with RNA data. Expression levels were provided as variance stabilization-transformed read counts from the DESeq package [32]. We further standardized RNA expression data for each gene by linearly scaling values across cell lines to a range between 0 and 1 and shifting the scaled values by subtracting the scaled mean. Gene identifiers in the protein-coding table were converted from Entrez Gene ID to ENSEMBL format using the org.Hs.eg.db package [33] in R, while gene identifiers in the non-coding table were already provided in ENSEMBL format. Once gene identifiers for the Klijn 2015 protein-coding and non-coding datasets were both in ENSEMBL format, we removed genes from the non-coding dataset that were found in the coding dataset. Following processing described above, the coding and non-coding expression data were merged and treated as a single dataset for downstream analysis.

For CCLE, we retrieved expression data from RNA-seq analysis as a single table from the CCLE database (file name: “CCLE_RNAseq_genes_rpkm_20180929.txt”; database file date: 29 September 2018). Expression levels for 55,414 genes (after excluding genes that were invariable in expression across all 1,019 CCLE cell lines) were provided as reads per kilobase of transcript per million mapped reads (RPKM). The natural log of CCLE RNA values was taken by log(RPKM+0.001), as the data exhibited a long right tail with a small number of extremely high expression values. CCLE gene identifiers for RNA expression data were provided in ENSEMBL format. CCLE RNA expression data was standardized as described above for the Klijn 2015 dataset (i.e. scaling and shifting). Standardization was performed independently for both Klijn 2015 and CCLE datasets. For downstream analysis (e.g. feature selection, model building), we included only the 31,047 genes with RNA expression data present in both Klijn 2015 and CCLE RNA-seq datasets, while genes present in only one dataset or the other were excluded.

### DNA variant data

For Klijn 2015, we retrieved single nucleotide polymorphism (SNP) and copy number variation (CNV) mutation data from the associated supplementary material (SNPs: “Supplementary Data 3” in [11]; CNVs: “Supplementary Data 4” in [11]). Klijn 2015 SNP and CNV data were identified from SNP microarray analysis (Illumina HumanOmni2.5_4v1). The provided SNPs had been previously filtered to include only missense and nonsense mutations. In all, there were 127,579 unique SNPs across 14,375 genes for all 675 cell lines with SNP calls. Because any given SNP was rare in the Klijn 2015 dataset, SNPs were mapped up to the gene level; genes then served as binary features (i.e. gene mutated / gene not mutated) without regard to which SNP(s) a gene contained. For Klijn 2015, CNV data were provided on a gene-by-gene basis as ploidy-adjusted copy number calls from the SNP array data using the PICNIC [34] and GISTIC [35] algorithms. We then classified the provided ploidy-adjusted copy number calls as amplifications (coded as 1) or deletions (coded as −1) based on thresholds described by Klijn et al [11] (amplifications: ≥1; deletions: ≤−0.75). In total, 18,396 gene amplifications and 42,371 gene deletions for 11,864 unique genes were present across 668 cell lines with CNV calls. Gene identifiers for both the SNP and CNV data in Klijn 2015 were converted from Entrez Gene ID to ENSEMBL format using the org.Hs.eg.db package [33].

For CCLE, we retrieved whole exome sequencing variant calling data from the CCLE database (file name: “CCLE_mutation_DepMap_18q3_maf_20180718.txt”; database file date: 18 July 2018). Of the 1,203,976 total variant calls present, only the 705,056 single-nucleotide variants annotated as either missense or nonsense mutations were included in further analysis – consistent with the SNPs available in the Klijn 2015 dataset. As described above for the Klijn 2015 dataset, SNPs were mapped up to the gene level where the mutation status of a gene served as a binary feature. Gene identifiers were converted from Entrez Gene ID to ENSEMBL format using the org.Hs.eg.db package [33]. CCLE CNV data were downloaded from the CCLE database (file name:

“CCLE_MUT_CNA_AMP_DEL_binary_Revealer.gct”; database file date: 29 February 2016). CNVs were provided based on calls from SNP microarray data (Affymetrix Genome-Wide Human SNP Array 6.0) and the GISTIC algorithm [35]. The downloaded CNV file provided gene-based amplification and deletion calls, which were assembled into a gene-by-cell line matrix with amplifications coded as 1, deletions coded as −1, and no copy number variation coded as 0. In total, 691,866 gene amplifications and 948,159 gene deletions were called across 1,030 cell lines. Gene symbols in the CCLE CNV data were converted to ENSEMBL IDs using the org.Hs.eg.db package [33]. In total, 12,399 genes mutated by SNP and 4,511 genes with CNVs were in common between the two datasets and included for downstream analysis. This analysis did not attempt to distinguish germline variants from somatic variants, mainly because there were no genotype data available to serve as the matched “normal” for the cell lines.

### Principal component analysis

We performed dimensionality reduction by principal component analysis (PCA) on a subset of genes with highly variable expression patterns. For each gene, the variance and mean were calculated for distributions of the pre-logged and pre-standardized expression values. A measure of expression variability was then calculated as the log of the ratio of variance over mean (values calculated in the previous step). We ranked expression variability of each gene based on log variance-to-mean ratio and included the union of the 3,000 most variable genes from the Klijn 2015 and CCLE datasets in dimensionality reduction analysis shown in Figure 1E.

### Cigar plots

We developed a visualization strategy to simultaneously display within- and between-group prediction performances, called the “cigar plot” (**Figure 3**). The cigar plot displays the predicted and observed drug response in an x-y plot, and highlights their concordance (i.e., the prediction performance) at two levels. First, the variation within each group is shown as an ellipse, with the within-group concordance (e.g., measured by rank correlation coefficients) shown as the tilt and width of the ellipses. Second, the between-group variation (differences of group mean values) is shown as the location of ellipsis centers. Ellipsis tilt was calculated by 45° × correlation coefficient: coefficient of 1 is associated with a 45° tilt, coefficient of 0 a 0° tilt, and coefficient of −1 a −45° tilt. The slope of the line across an ellipse is the same as ellipsis tilt. In **Figure 3**, rank correlation coefficients (Spearman’s ρ) were used. Ellipsis length is scaled to the observed within-group range, assessed on a per-tissue basis as by concatenating the 4 sets of MEKi responses in both datasets and calculating the interquartile range across all MEKi response values. Ellipsis width scales with within-group performance and is calculated as: ellipsis length × (1 - absolute correlation coefficient). Larger absolute coefficients result in narrower ellipses (coefficient of 1 results in a line) while smaller coefficients result in rounder ellipses (coefficient of 0 results in a circle). R code to generate cigar plots is available at: https://github.com/johnplloyd/cigar_plot.

### Model building approach

We developed prediction models for four sets of input data (“4 models” in **Figure 2A-B**), trained by considering drug response data for the two MEK inhibitors in the two datasets as the outcome, with the associated molecular features as the independent variables– e.g. the response to PD-901 in Klijn 2015, to be predicted with the Klijn 2015 molecular features (**Figure 2A**). The two models from Klijn 2015 were denoted *f_K1_*, *f_K2_*, likewise the two from CCLE were denoted *f_C1_*, *f_C2_*. Within each model, we applied four algorithms: regularized regression, random forest (regression), logistic regression, and random forest (classification) prediction models (additional details below). In all, we trained 4 MEKi × 4 algorithms = 16 prediction models. To assess performance, we performed two types of validation: 1) within-dataset validation: training a model in a dataset using the non-shared cell lines (i.e. unique to a given dataset), predicting the drug response for the other, 154 shared cell lines using their molecular features in this same dataset, and compared with their actual drug response; 2) between-dataset validation: training a model in the same way as above (i.e., using the non-shared cell lines), predicting the drug response for all cell lines (both shared and non-shared) in the other dataset, using the molecular features in the other dataset, and compare with the actual drug response in the other dataset. An example schematic of model training while considering PD-901 in the Klijn 2015 dataset and model application both within and across datasets is shown in **Figure 2A**.

During the training phase, for regression-based algorithms, 30 prediction models were built after randomly-sampling 70% of available training cell lines, and the predicted validation set response was calculated as the average of the 30 randomly-sampled runs. This procedure is conceptually similar to a bootstrap aggregating (bagging) approach, but differs in that each of the 30 random training sets was sampled *without* replacement (whereas bagging methods are defined by sampling *with* replacement). Note that the remaining 30% of training cell lines that were not selected for each of the 30 iterations of model training were not used to assess model performance. For classification-based algorithms, 30 prediction models were trained using an equal number of sensitive and resistant cell lines: 70% of cell lines of the least populated class (typically sensitive) in the training set were randomly selected and matched with an equal number of randomly-selected cell lines of the more populated class (typically resistant). As with regression-based algorithms, classification-based prediction models were applied to within-dataset and cross-dataset validation sets and the final prediction score was calculated as the average of 30 prediction models.

As mentioned above, we established sets of regularized regression, random forest (regression), logistic regression, and random forest (classification) prediction models. Regularized regression and random forest (for both regression and classification) models were trained using the *glmnet* [36] and *randomForest* [37] packages in R, respectively. Logistic regression models were trained using the lm() function (family = binomial) in the base installation of R. Regression prediction models (regularized and random forest) were trained on log(IC_50_) of MEKi response. Classification prediction models (logistic and random forest) were trained on binarized IC_50_ MEKi response, with IC_50_ ≤1 defining sensitive cell lines and IC_50_ > 1 defining resistant.

For regularized regression predictions, the α parameter controls whether a ridge regression (α = 0), elastic net (0 < α < 1), or least absolute shrinkage and selection operator (LASSO; α = 1) model is generated and the λ parameter controls the strength of the penalty on model ß values. We tuned the α and λ parameters through a parameter sweep. We tested α values from 0 to 1 in 0.1 step increments, while the λ values tested were 0.001, 0.01, 0.1, 1, and 10. For logistic regression and random forest (regression and classification) models, we performed feature selection by LASSO prior to model training. For the LASSO feature selection, we performed a parameter sweep to test multiple λ values: 1×10^−5^, 5×10^−5^, 1×10^−4^, 5×10^−4^, 0.001, 0.005, 0.01, 0.05, and 0.1. Tuning of α and λ parameters was performed by randomly selecting 70% of training cell lines and applying models trained with different parameter sets to the remaining 30% of training cell lines, repeated for 100 iterations. Note that the validation cell lines were not used for parameter tuning. Optimal parameter sets were selected based on maximum mean performance in the testing set across the 100 iterations. Performance was measured by Spearman’s ρ for regression algorithms and auROC for classification algorithms. **Table S1** shows the optimal parameters selected.

For tissue-specific models, we trained the prediction models with CCLE data, randomly selecting 75% of cell lines of a given cancer type and applied the models to Klijn 2015 data for testing. If a cell line overlapping between Klijn 2015 and CCLE was selected to build the tissue-specific model, it was excluded from model testing. Tissue-specific prediction models were the only instance in which the 154 cell lines in common between Klijn 2015 and CCLE were used for training, as there were fewer cell lines available for individual cancer types. Parameters selected from pan-cancer prediction models (**Table S1**) were also used for generating tissue-specific models.

## ACKNOWLEDGEMENTS

Work was supported by funds from the Michigan Institute for Clinical and Health Research – Postdoctoral Translational Scholar Program (michr.umich.edu), Breast Cancer Research Foundation (brcf.org), Michigan Institute for Data Science (midas.umich.edu), and National Institutes of Health (nih.gov; NIH 1R21CA218498-01 to S.D.M. and NIH R01GM118928-01 to J.Z.L.). The funders had no role in study design, data collection and analysis, decision to publish, or preparation of the manuscript.

## AUTHOR CONTRIBUTIONS

**Conceptualization:** JPL, MS, SDM, JZL

**Formal Analysis:** JPL

**Funding Acquisition:** JPL, SDM, JZL

**Writing – Original Draft Preparation:** JPL

**Writing – Review & Editing:** JPL, MS, SDM, JZL

## DECLARATION OF INTERESTS

The authors have declared that no competing interests exist.

## SUPPLEMENTAL LEGENDS

**Supplemental Figure 1. Within- and between-tissue RNA expression similarity.** Heatmap colors indicate the mean ranked RNA expression correlation (Spearman’s ρ) across highly variable genes (see Methods) for 100 randomly-selected pairs of cell lines of the indicated tissues. Black outlines indicate comparisons within the same tissue type. Expression correlation is shown for cell lines **(A)** within Klijn 2015, **(B)** within CCLE, and **(C)** between the two datasets.

**Supplemental Figure 2. Prediction performances as assessed in the full pan-cancer cell line set (left-most column) and within 10 cancer types (the next 10 columns).** Heatmaps indicate rank correlation (Spearman’s ρ) between observed and predicted MEKi responses (**Figure 2**) based on regularized (top) and logistic (bottom) regression prediction models. Each row is for a specific combination of training data and test data, over two MEK inhibitors and two datasets. Also shown are the mean performance for each column (bottom row). Dendrogram at the top depicts hierarchical clustering of the tissues by their performance patterns.

**Supplemental Figure 3. Within- and between-tissue signals for the top 50 marker genes based on their maximum absolute regularized regression coefficient.** (Left) Probability density distributions for a measure of within-tissue signal: the % of within-tissue variance in MEKi response explained for the top 50 markers (red) and all genes (gray). (Right) Probability density distributions for a measure of between-tissue signal: the absolute correlation (*r*) between the mean per-tissue gene expression and the mean per-tissue MEKi response. Vertical dotted lines indicate median values for color-matched distributions. *P*-values are from Mann-Whitney U tests comparing the two distributions in each panel. **(A-D)** Results are shown for the four MEKi screens.

**Supplemental Table 1. Optimal model parameters.** Model parameters (λ and α) selected under cross-validation within training sets.

## Notes

### Competing Interest Statement

The authors have declared no competing interest.

