## Supplementary Information for "Impact of between-tissue differences on pan-cancer predictions of drug sensitivity"

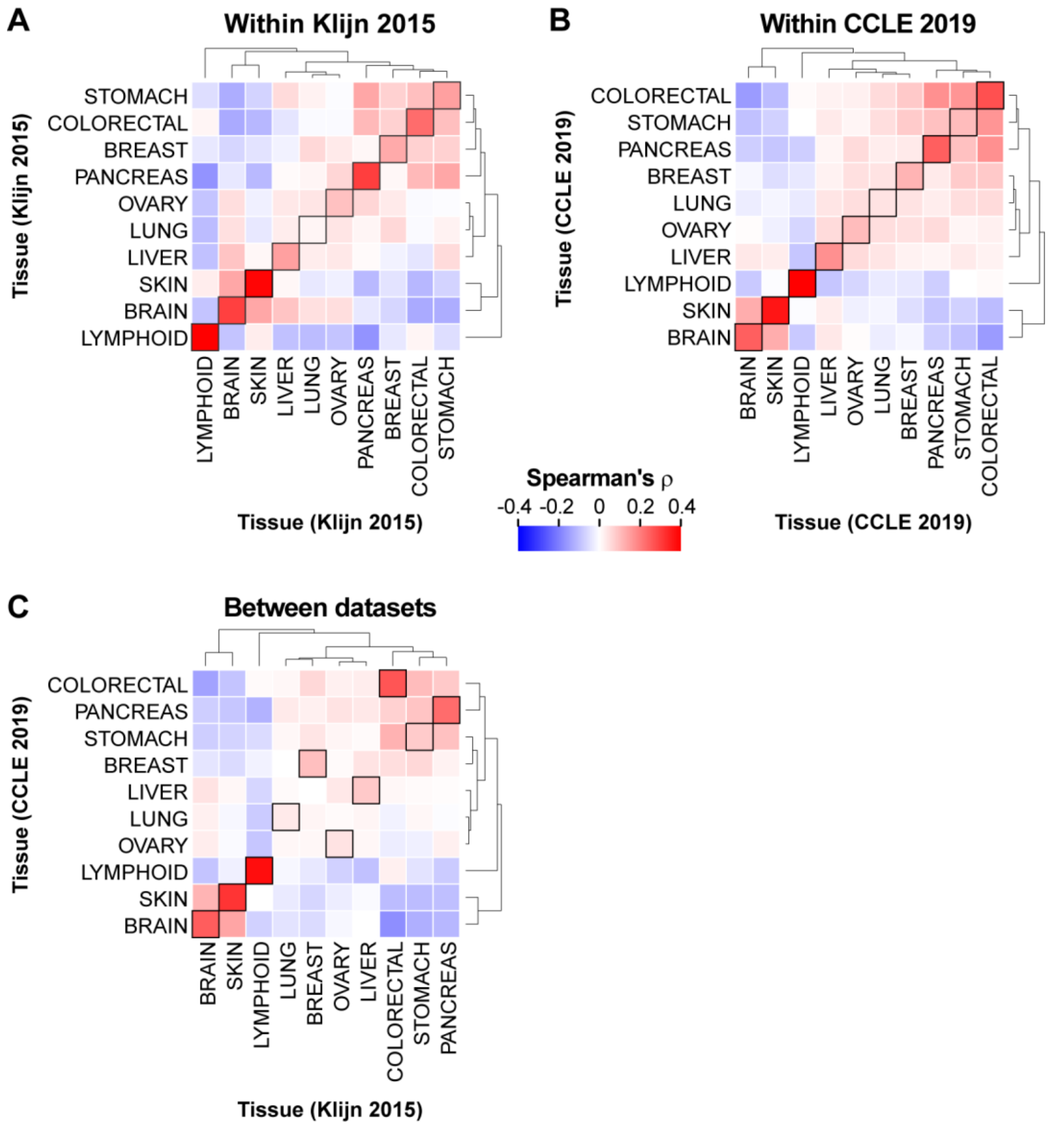


**Supplementary Figure 1.** Within- and between-tissue RNA expression similarity. Heatmap colors indicate the mean ranked RNA expression correlation (Spearman's ρ) across highly variable genes (see Methods) for 100 randomly-selected pairs of cell lines of the indicated tissues. Black outlines indicate comparisons within the same tissue type. Expression correlation is shown for cell lines **(A)** within Klijn 2015, **(B)** within CCLE, and **(C)** between the two datasets.

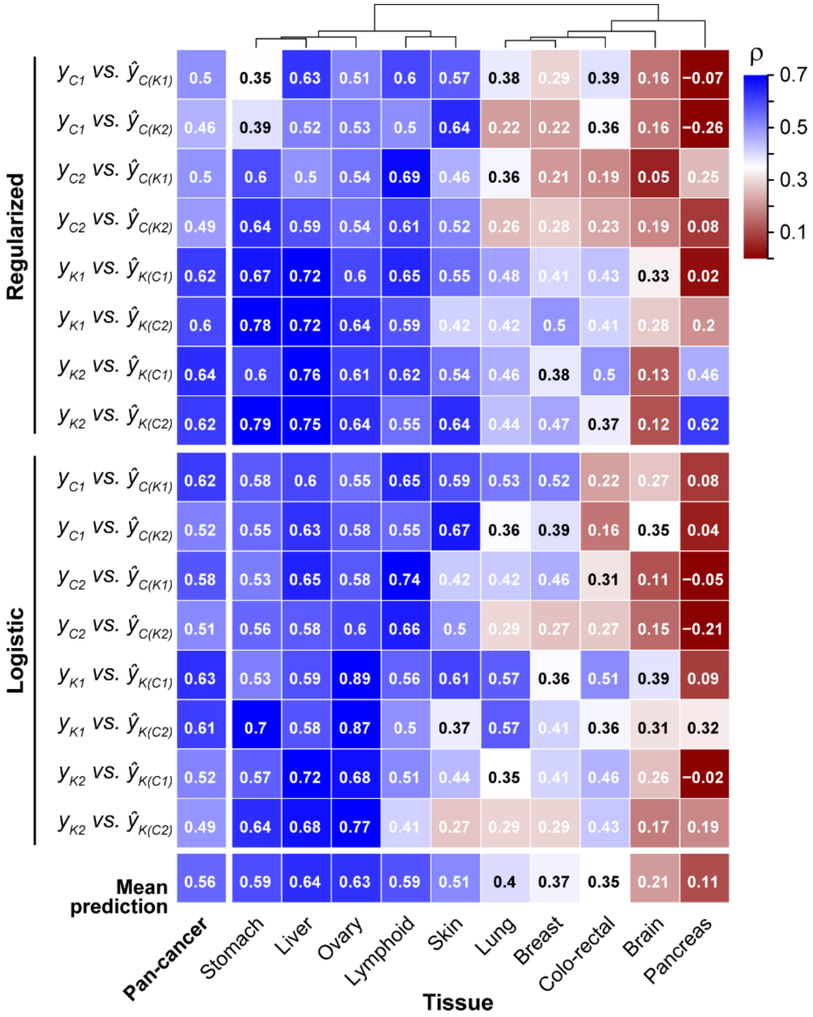


**Supplementary Figure 2.** Prediction performances as assessed in the full pan-cancer cell line set (left-most column) and within 10 cancer types (the next 10 columns). Heatmaps indicate rank correlation (Spearman's ρ) between observed and predicted MEKi responses (**Figure 2**) based on regularized (top) and logistic (bottom) regression prediction models. Each row is for a specific combination of training data and test data, over two MEK inhibitors and two datasets. Also shown are the mean performance for each column (bottom row). Dendrogram at the top depicts hierarchical clustering of the tissues by their performance patterns.

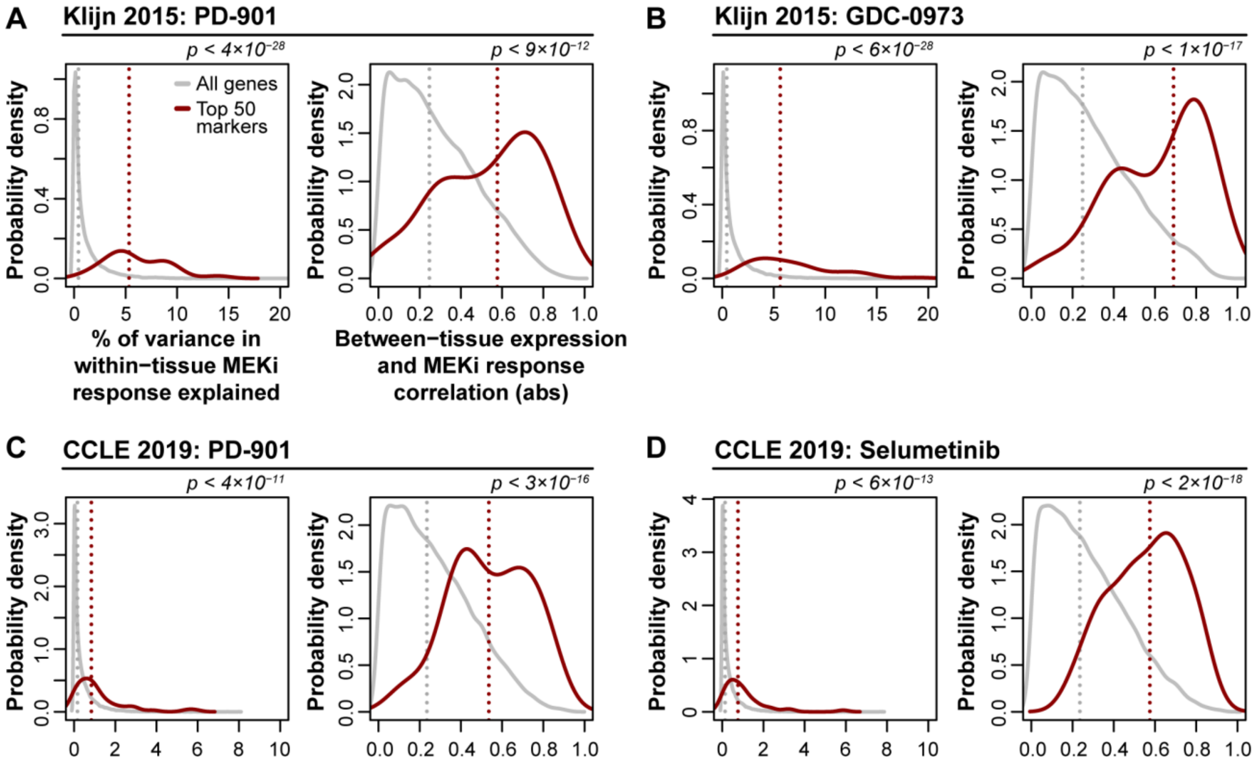


**Supplementary Figure 3.** Within- and between-tissue signals for the top 50 marker genes based on their maximum absolute regularized regression coefficient. (Left) Probability density distributions for a measure of within-tissue signal: the % of within-tissue variance in MEKi response explained for the top 50 markers (red) and all genes (gray). (Right) Probability density distributions for a measure of between-tissue signal: the absolute correlation (*r*) between the mean per-tissue gene expression and the mean per-tissue MEKi response. Vertical dotted lines indicate median values for color-matched distributions. *P*-values are from Mann-Whitney U tests comparing the two distributions in each panel. **(A-D)** Results are shown for the four MEKi screens.

**Supplementary Table 1. Optimal model parameters.**

| Algorithm | Model | $\lambda$ [1] | $\alpha$ [2] |
|---|---|---|---|
| Regularized regression | $f_{K1}$ | 1 | 0.1 |
| | $f_{K2}$ | 1 | 0.1 |
| | $f_{C1}$ | 10 | 0 |
| | $f_{C2}$ | 10 | 0 |
| Random forest regression | $f_{K1}$ | 0 (no feature selection) | *NA* |
| | $f_{K2}$ | 0 (no feature selection) | *NA* |
| | $f_{C1}$ | 0 (no feature selection) | *NA* |
| | $f_{C2}$ | 0 (no feature selection) | *NA* |
| Logistic regression | $f_{K1}$ | 0.05 | *NA* |
| | $f_{K2}$ | 0.1 | *NA* |
| | $f_{C1}$ | 0.01 | *NA* |
| | $f_{C2}$ | 0.1 | *NA* |
| Binary random forest | $f_{K1}$ | $1x10^{-5}$ | *NA* |
| | $f_{K2}$ | $1x10^{-5}$ | *NA* |
| | $f_{C1}$ | $1x10^{-5}$ | *NA* |
| | $f_{C2}$ | $1x10^{-5}$ | *NA* |

[1] For regularized regression λ was used for both feature selection and β-penalization. For logistic regression and both random forest methods, λ was used for feature selection only.

[2] α was used to evaluate three regularization methods: ridge regression (α = 0), elastic net (0 < α < 1), and LASSO (α = 1). *NA*: not applicable, α is not a parameter for the non-regularized algorithms.
